# Comparative analysis of Formyl peptide receptor 1 and Formyl peptide receptor 2 reveals shared and preserved signaling profiles

**DOI:** 10.1101/2024.02.08.579483

**Authors:** Denise Pajonczyk, Merle F. Sternschulte, Oliver Soehnlein, Marcel Bermudez, Carsten A. Raabe, Ursula Rescher

## Abstract

Pattern Recognition Receptors are key in identifying pathogenic or damaged cell-related patterns or molecules. Among these, the closely linked formyl peptide receptors FPR1 and FPR2 are believed to hold pivotal yet differing functions in immune regulation. To address the intriguing question of how these highly related receptors with a shared agonist spectrum play differing roles in modulating inflammation, we analyzed the signaling profile for a panel of FPR agonists *in vivo* and *ex vivo* settings. Our analysis uncovered a shared core signature for both FPRs across signaling pathways. Whereas formylated peptides generally acted as potent agonists at FPR1, FPR2 agonists, irrespective of N-terminal formylation, displayed consistently low activity ratios, suggesting an underutilized signaling potential of this receptor. Signaling outcomes were defined by specific agonist-receptor pairings and no receptor-specific signaling texture was identified. Activation of the FPR signaling axis by fMLF in human neutrophils did impact neutrophil survival. Overall, the distinct characteristics underlying inflammatory, anti-inflammatory, or pro-resolving profiles could not be attributed to a specific receptor isoform, signaling pattern, or a particular class of agonists, challenging assumptions about distinct inflammatory profiles linked to specific receptors, signaling patterns, or agonist classes.

## Introduction

The immune system relies on Pattern Recognition Receptors (PRRs) to detect specific patterns or molecules associated with threats like pathogens or cellular damage [1]. Activated PRRs initiate the innate immune defense by recognizing Pathogen-Associated Molecular Patterns (PAMPs) found in pathogens but not in the host, and Danger-Associated Molecular Patterns (DAMPs) released by damaged cells [2], [3].

Bacterial but not eukaryotic protein translation is initiated by N-formylated methionine. The modified amino acid, therefore, serves as a PAMP [4]–[6]. Formyl Peptide Receptor 1 (FPR1) was identified as the cognate receptor for the bacterial chemotactic and pro-inflammatory tripeptide fMLF (formyl-Met-Leu-Phe)[7]–[9]. Evolutionary analysis revealed FPR1 as the founding member of the FPR family, which are PRRs that belong to the seven transmembrane G protein-coupled receptors (GPCRs) [6], [10], [11] and are predominantly expressed on immune cells [12], [13]. Mitochondrial protein synthesis, like bacteria, uses N-formylated methionine for initiation [14]–[17]. Consequently, FPRs also detect peptides released from damaged mitochondria and thus integrate the immune response to bacterial infection and/or tissue degradation [6], [13]. Notably, the repertoire of FPR ligands has significantly expanded beyond the recognition of only formylated DAMPs and PAMPs. FPR ligands are highly diverse and include non-formylated peptides of viral and bacterial origins, host-derived polypeptides, and even lipids [8], and often target both FPR1 and the closely related FPR2 [6], [18]. Apart from these naturally occurring FPR ligands, the potent pan-FPR agonist W-peptide (WKYMVm) and the FPR2-selective agonist MMK 1 (LESIFRSLLFRVM) were the results of artificial peptide synthesis and library screening [19]–[22]. FPRs were originally believed to mediate pro-inflammatory responses. In various progressive disease states, however, FPR2 knockout models revealed the involvement of FPR2 signaling to the resolution of inflammation [23]. Lack of FPR2 signaling has been linked to exacerbated injury in animal models of ischemia-reperfusion injury, associated with neutrophil pro-inflammatory activation [24]. In particular, annexin A1, a protein also abundantly expressed in neutrophils [25], has been linked to the regulation of inflammation and the resolution of inflammatory processes via FPR2 activation [26]–[28]. The anti-inflammatory and pro-resolving abilities induced by FPR2 agonists have positioned this receptor as a promising target for drug development This, in particular, is being enhanced by recent structural analysis of ligand-receptor interactions on the atomic scale [29]–[31]. FPR1 on the other hand has not been assigned anti-inflammatory roles so far; notably, both receptors share an amino acid sequence identity of approximately 69% and hence display a high degree structural similarity ([6], [32]). Given their homology, structural relation and shared agonist repertoire, including annexin A1 and its mimetic N-terminal peptide Ac2-26 [6], [26], [33], [34], we explored whether and how these closely related receptors with overlapping expression patterns [6], [34] convey distinct signaling profiles. Potentially, each receptor might have a distinct signaling profile or recognize FPR agonist subsets with specific activation patterns [35]. For analysis, we compared a set of representative FPR peptide agonists encompassing host- and pathogen-derived as well as synthetic ligands across cellular pathways linked to GPCRs — such as Gi-mediated inhibition of de novo cAMP generation, activation of the MAPKinase pathway, and receptor internalization [36], [37].

Our comprehensive analysis of agonist potencies (apparent EC50), apparent efficacies (Emax), and activity ratios, which integrates both logistic parameters, revealed a shared core signature for both FPR1 and FPR2. Notably, the synthetic agonist W-peptide exhibited the strongest agonism both in terms of potency and efficacy at both receptors and across all pathways, defining the maximum activity ratio and maximum space of receptor activation in our settings. Confirming our previous report [37], shorter formylated peptides triggered FPR1 activity to the maximum. In stark contrast, the activity ratios of FPR2 agonists, regardless of their origin, were consistently lower than those induced by W-peptide, suggesting that endogenous agonists do not fully exploit the entire signaling potential of FPR2. Overall, signaling outcomes were not linked to a particular receptor isoform, signaling texture, or a specific class of agonists, such as formylated peptides. Rather, they were dependent on the specific pairing of agonist and receptor.

FPR1 and FPR2 are robustly expressed in human neutrophils [6], [38]. In circulation, these non-dividing cells are generally relatively short-lived, yet their lifespan can extend considerably in the presence of inflammatory signals including endotoxins to increase neutrophil-mediated innate immune defense [39], [40]. The reversal of this initially beneficial reaction by induction of apoptosis is a crucial part of pro-resolving mechanisms as it prompts the removal of neutrophils and limits excessive tissue damage caused by prolonged neutrophil activity [39], [41], [42]. Here, neither the pan agonist WKYMVm nor the ligands fMLF and MMK 1 could reverse the LPS-induced prolonged neutrophil survival. However, fMLF treatment per se displayed significant lower numbers of apoptotic neutrophils, agreeing with the idea of FPR1 and its formylated ligands as pro-inflammatory mediators [37], [43] .

## Results

### G*α*i-dependent activation patterns at FPR1 and FPR2 are similar

FPR1 and FPR2 share are conserved three-dimensionally structure. While the full-length sequence identity is 68.6%, the structure-determined parts show 70.5% identity and 82.2% similarity, and binding site residues display an even higher identity of 73.1% and similarity of 88.5% (Fig. 1), underlining the close functional relationship between the two receptors.

**Figure 1.**
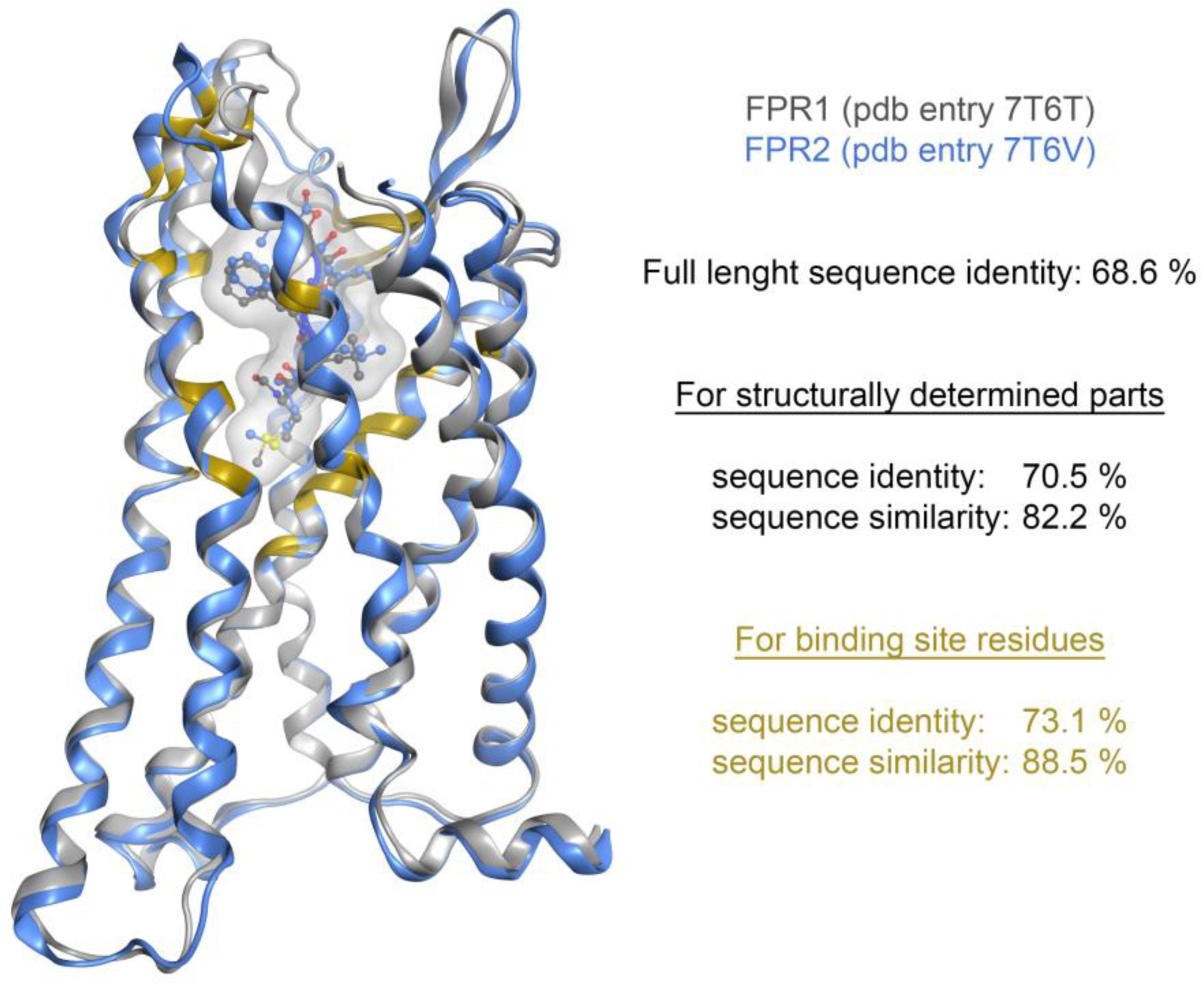
Structural Comparison of Human Formyl Peptide Receptors. The structural overlay illustrates the three-dimensional similarities and differences between FPR1 (pdb entry 7T6T, grey ribbon) and FPR2 (pdb entry 7T6V, blue ribbon), emphasizing their conserved structure with high sequence identity and similarity. The structural alignment and visualization was performed with MOE (Molecular Operating Environment, 2022.02, Chemical Computing Group ULC).

Despite their high homology, FPR1 and FPR2 are reported to exert distinct functions in the immune response [36]. For the comparison of ligand-induced signaling profiles, we generated HEK293 cell lines ectopically expressing either FLAG-tagged FPR1 or FPR2 (for analysis of the receptor expression and correct plasma membrane localization, see suppl. Fig S1). The parental cell line was not responsive to the FPR ligands (suppl. Fig. S2). We recorded concentration-response curves for the inhibition of *de novo* cAMP formation and ERK/MAPK pathway activation for a panel of PAMP and DAMP agonists that interact with FPR1 (Fig. 2A) and FPR2 (Fig. 2B). The test systems were validated with the prototypic FPR1 agonist fMLF, the FPR2-specific agonist MMK 1[19], [44], and the synthetic pan-FPR agonist W-peptide [8], [21], [22]. As expected, W-peptide featured as a highly potent agonist, showing nanomolar EC50 values for both receptors and pathways. MMK 1, on the other hand, exclusively activated FPR2-dependent signaling, while fMLF triggered FPR1-mediated signaling in the nanomolar range but was less potent on FPR2. Formylated PAMPs and DAMPs activated either receptor, though to varying extents. Notably, MT-CytB did not elicit detectable ERK1/2 activation at FPR1. To explore if host-derived non-formylated FPR agonists trigger a distinct signaling profile compared to formylated peptides, we introduced mitochondria-derived peptides encoded within the 16S rRNA gene—specifically humanin and the small humanin-like peptide 6 (SHLP6) [45]–[47]. Humanin, originally described as exclusive FPR2 agonist [48] with defined anti-inflammatory properties, surprisingly also induced FPR1 activity. In contrast, SHLP6 triggered ERK1/2 activation solely at FPR1. Additionally, we included the annexin peptide Ac2-26, a recognized anti-inflammatory mediator acting via FPRs [49], [50].

**Figure 2.**
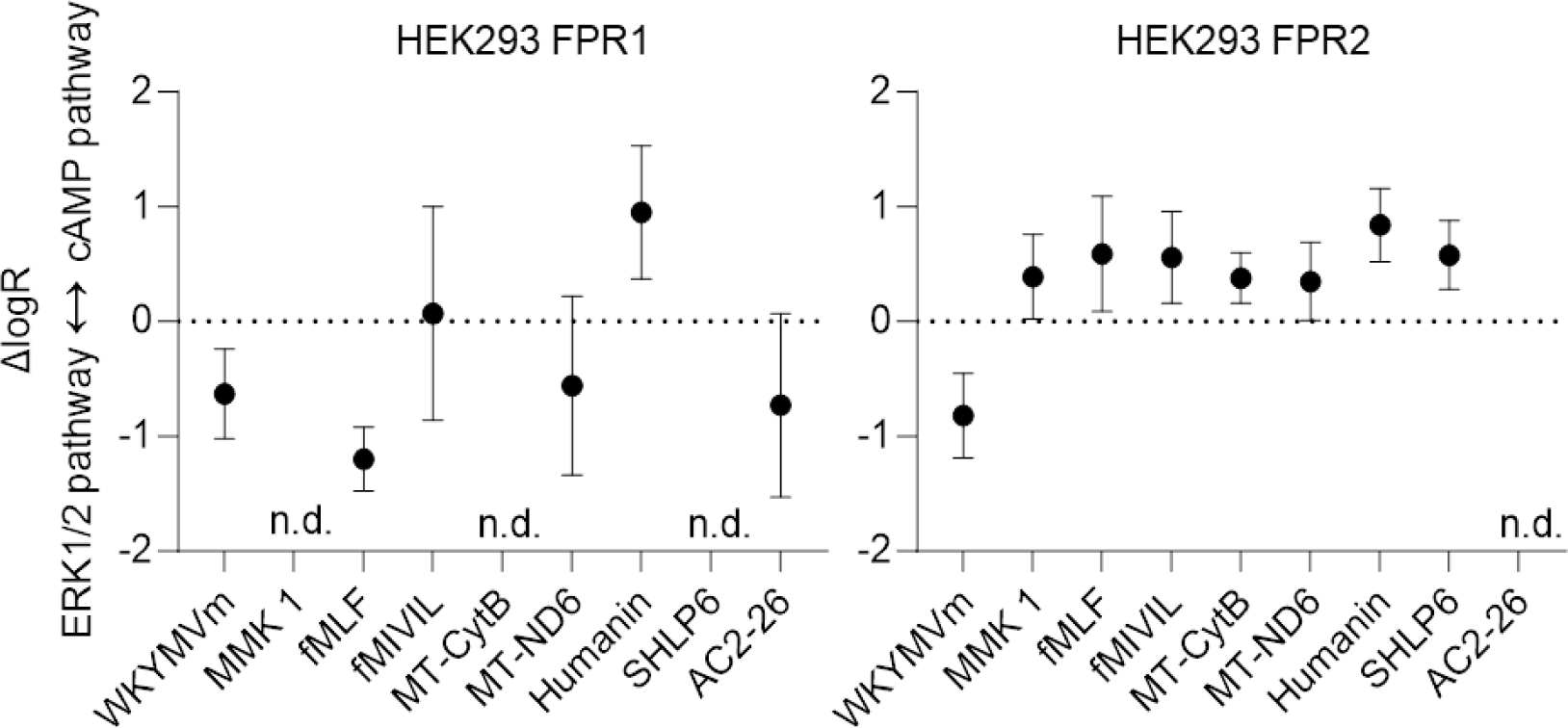
Pathway preferences. To assess the preferred signaling pathway, ΔlogRpathway values were calculated as the difference of logRERK1/2activation - logRinhibitionofcAMP formation. Positive values indicate a preference for ERK1/2 activation; negative values indicate that the inhibition of de novo cAMP formation is favored. n.d.: not detectable.

In agreement with our previous observations [50], Ac2-26 acted as an agonist at both receptors, with a preference for FPR1. However, the peptide failed to inhibit *de novo* cAMP generation at FPR2, similar to SHLP6 at FPR1. No distinct signaling pattern could be attributed to either the synthetic, formylated PAMPs and DAMPs, or non-formylated host-derived agonists. As anticipated [37], activation profiles of human FPR1 and FPR2 assessed in the presence of pertussis toxin (PTX) confirmed the dependence on Gαi-mediated signaling ([51] & Suppl. Fig. 2).

An overview of the logistic parameters for both receptors across pathways is shown in suppl. Fig. 3 and in Table I. We next deduced logR values, the logarithm of the ratio of both the logistic parameters EC50 and the Emax value of the respective agonist and pathway [52]. To assess agonist-specific pathway preferences, we computed ΔlogR_pathway_ values by subtracting logR values for ERK1/2 activation from those for *de novo* cAMP formation. No specific agonist group was associated with preferential pathway activation (Fig. 3). Instead; the activation preferences were contingent on the specific combination of receptor and agonist, which was particularly evident for FPR1. At FPR2, all physiological agonists displayed a preference toward inhibition of *de novo* cAMP generation. Notably, however, FPR2 activation by W-peptide revealed the potential of the receptor to preferentially activate ERK1/2.

**Figure 1.**
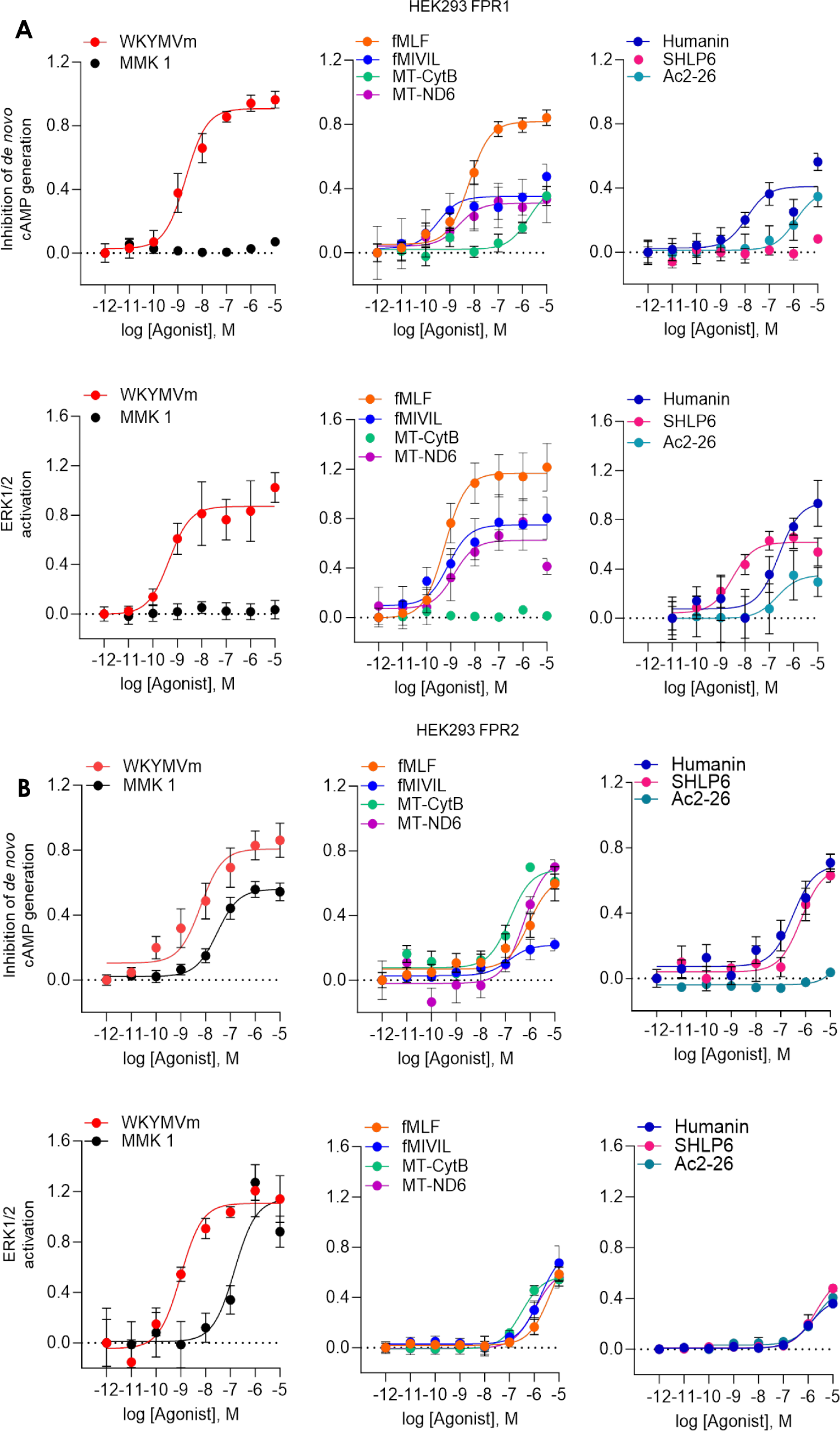
Activation profiles of human FPR1 and FPR2. For analysis of the inhibition of *de novo* cAMP formation, HEK293 FPR1 (A) and FPR2 (B) cells were stimulated with forskolin and the indicated FPR agonist, and responses were recorded 30 min after agonist addition. ERK1/2 activation was monitored 5 min after the addition of the agonist. Concentration-response curves were fitted using an LL4 model. Data points display means ± SEM of n = 5 or 6 independent experiments.

W-peptide featured consistently as the agonist with the highest activity ratio at both receptors and for both pathways. For compound comparisons, we selected this maximum activator as the internal standard and normalized agonist logR values to this reference, resulting in ΔlogR_agonist_ values (Fig. 4). At FPR1, the bacteria-derived formylated peptides (fMLF, fMIVIL) showed maximum receptor activation similar to W-peptide in both pathways. The endogenous mitochondrial formylated peptide MT-ND6 was also classified among these maximum FPR1-activating agonists. However, MT-CytB presented a reduced or even absent induction of FPR1 activity, a finding that is in agreement with a preference for FPR1 for smaller formylated peptides like fMLF [6], [32]. At FPR2, the synthetic ligands MMK 1 and W-peptide displayed similar signaling profiles for inhibition of cAMP generation, yet MMK 1 was less potent than W-peptide for ERK1/2 activation.

**Figure 4.**
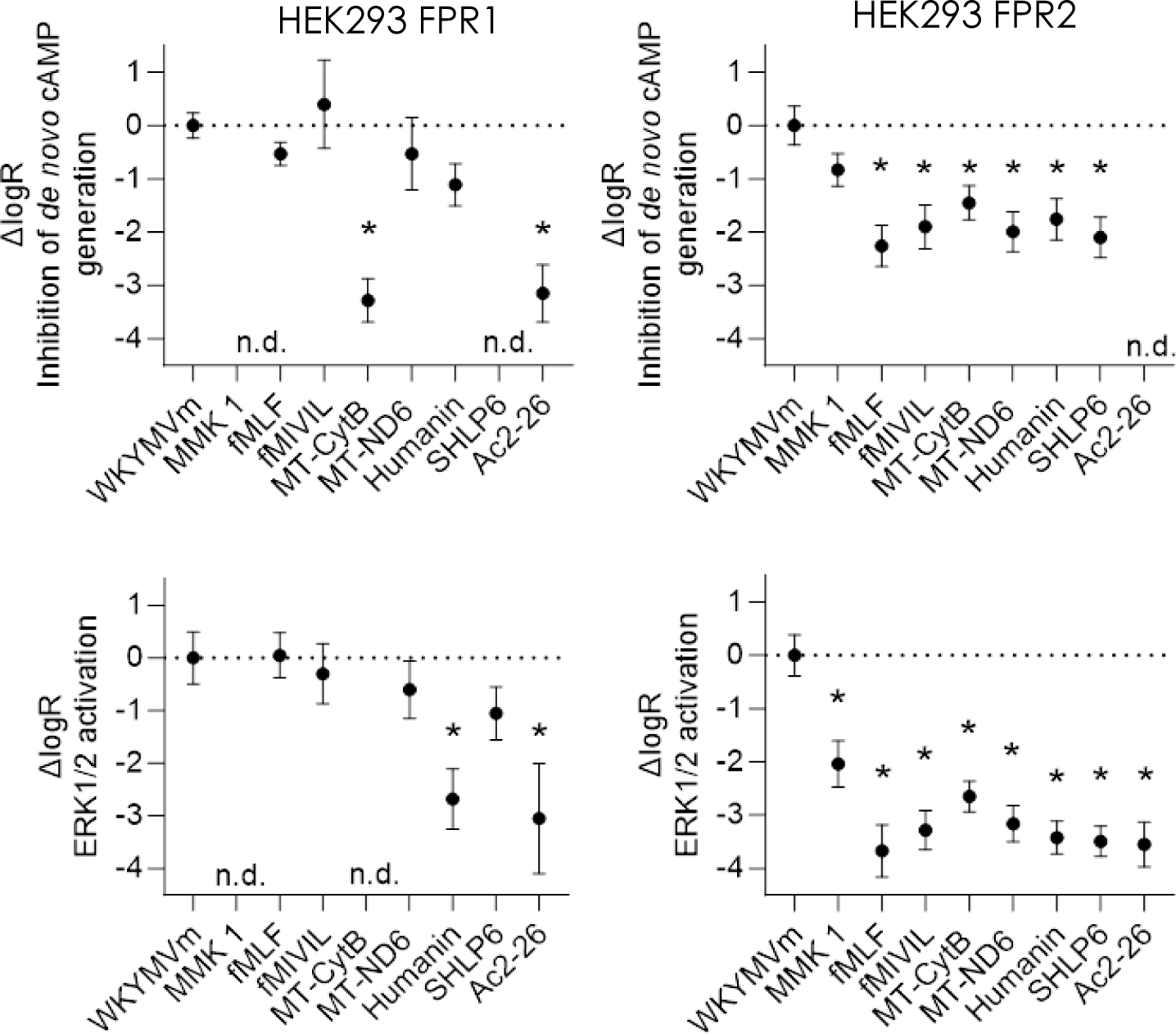
Comparison of agonist activities. ΔlogRagonist values with W-peptide as the reference agonist were calculated (n= 5-6). One-way ANOVA with Dunnett’s test; * p < 0.05 in comparison to W-peptide. n.d = not detectable.

The ΔlogR_agonist_ values of the remaining ligands, both formylated and non-formylated, differed significantly from W-peptide in both pathways. The efficient activation of FPR2 by W-peptide and MMK 1, suggested that natural agonists do not fully exploit the FPR2 signaling potential.

### Agonist-induced internalization of FPR1 and FPR2

We further investigated receptor internalization [53], a pathway less reliant on G protein engagement ([37] and suppl. Fig. 3). Again, W-peptide was the maximum activator at both receptors. Interestingly, only formylated peptides led to noticeable levels of receptor internalization (Fig. 5), reinforcing the evolutionary conserved preference of both receptors for this molecular pattern. However, the receptor preferences for shorter or longer peptides were evident by the activity ratios obtained for MT-CytB and fMLF. Formylated peptides triggered FPR1 internalization to levels achieved by W-peptide.

**Figure 5.**
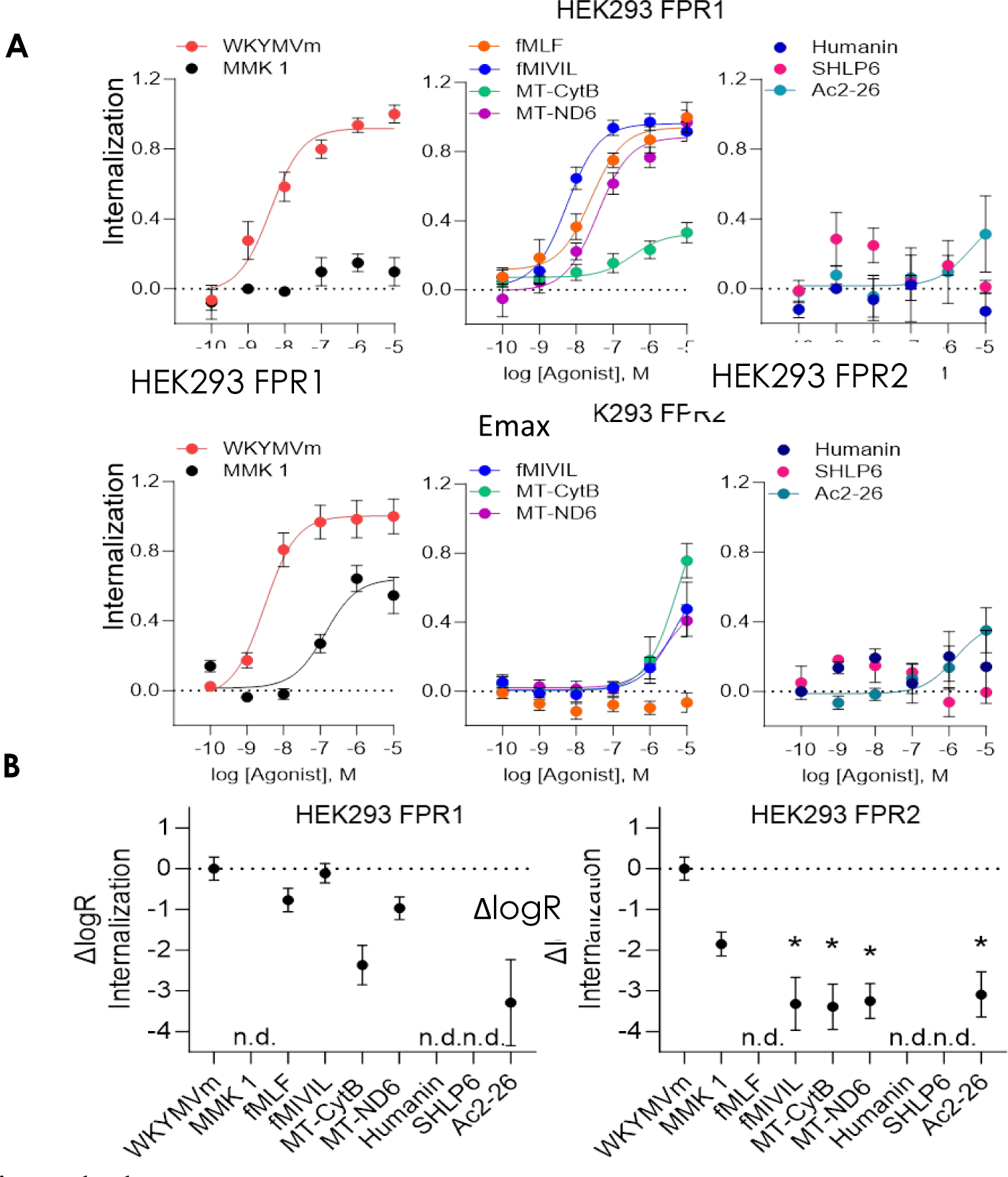
Agonist-induced receptor internalization profiles. (A) Concentration-response curves were recorded and fitted using an LL4 model. Data points indicate means ± SEM of 5 or 6 independent experiments. (B) ΔlogR values were calculated with W-peptide as the reference agonist, one-way ANOVA with Dunnett’s Test; * p < 0.05 in comparison to W-peptide. n.d. = not detectable

Conversely, no agonist, including MMK 1, induced FPR2 internalization to levels comparable to the reference agonist. Similar to the results for the Gαi-dependent pathway activation, the FPR2 signaling potential was again not fully exploited. Humanin and SHLP6 were not able to induce internalization of either receptor, suggesting that their signaling profiles were biased toward activation of G protein-dependent signaling. In contrast, Ac2-26 triggered the internalization of both FPR1 and FPR2, confirming its agonistic nature at FPR1 and FPR2. Fig. 6 summarizes the signaling profiles of the agonists at both receptors.

**Figure 6.**
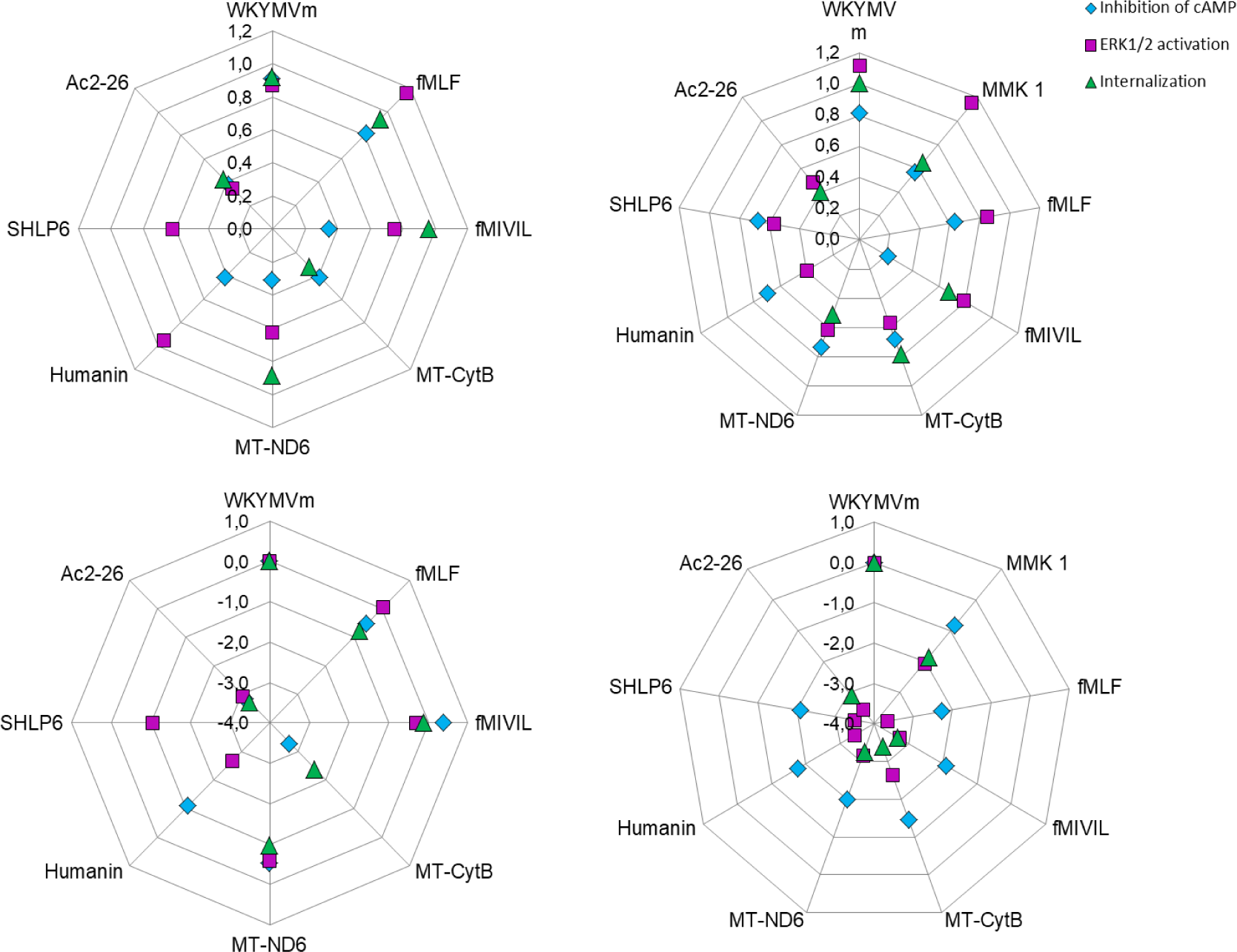
Graphical representation of the relative agonist activities. Each radius for ΔlogR is displayed in the logarithmic scale; radiuses for Emax are scaled linearly.

### Endotoxin-induced neutrophil long-term survival is not reversed by FPR1/2 activation

From our *in vitro* data, no distinct receptor-specific signaling profile could be assigned. Because FPR1 and FPR2 are robustly expressed in immune cells such as neutrophils [34], we next explored receptor activation in this physiological context to analyze whether signals derived from the receptor-specific engagement on these cells cause responses unique to either FPR1 or FPR2. The majority of circulating neutrophils are relatively short-lived and undergo spontaneous apoptosis within hours but inflammatory signals such as endotoxins prolong their lifespan considerably as part of the innate immune defense [42], [54], [55]. Hence, the reversal of increased neutrophil lifespan by induction of apoptosis is a crucial part of pro-resolving mechanisms as it prompts the removal of neutrophils and limits excessive tissue damage caused by prolonged neutrophil activity [40], [55].

We next addressed FPR activation in this model. A hallmark of early apoptosis is the translocation of phosphatidylserine (PS) from the inner cellular membrane to the outer leaflet cell surface (that can be monitored by binding of annexin V to PS), while the membrane remains impermeable [55], thus preventing binding of propidium iodide to DNA, allowing for the quantitation of early apoptotic (AnxV+/PI-) cells by flow cytometry. LPS-induced prolongation of neutrophil lifespan served as a positive control. To differentiate receptor subtype-specific effects, we tested fMLF and MMK 1 and also investigated the simultaneous activation of both receptors by utilizing the pan-agonist W-peptide.

A representative density dot plot is presented in suppl. Fig. S4. Approximately 60% of spontaneously apoptotic neutrophils were identified in the total cell population after 20 h (Fig. 7). As anticipated, LPS treatment increased neutrophil survival [41], leading to a significantly lower number of apoptotic neutrophil 20 hours of incubation. When compared to untreated neutrophils, fMLF treatment lowered the apoptotic cell count significantly. FPR activation with W-peptide slightly yet non-significantly reduced the number of apoptotic cells. None of the tested representative FPR agonists counteracted the LPS-induced neutrophil survival, suggesting that the FPR signaling increases neutrophil lifespan, in line with transduction of a pro-inflammatory stimulus.

**Figure 7.**
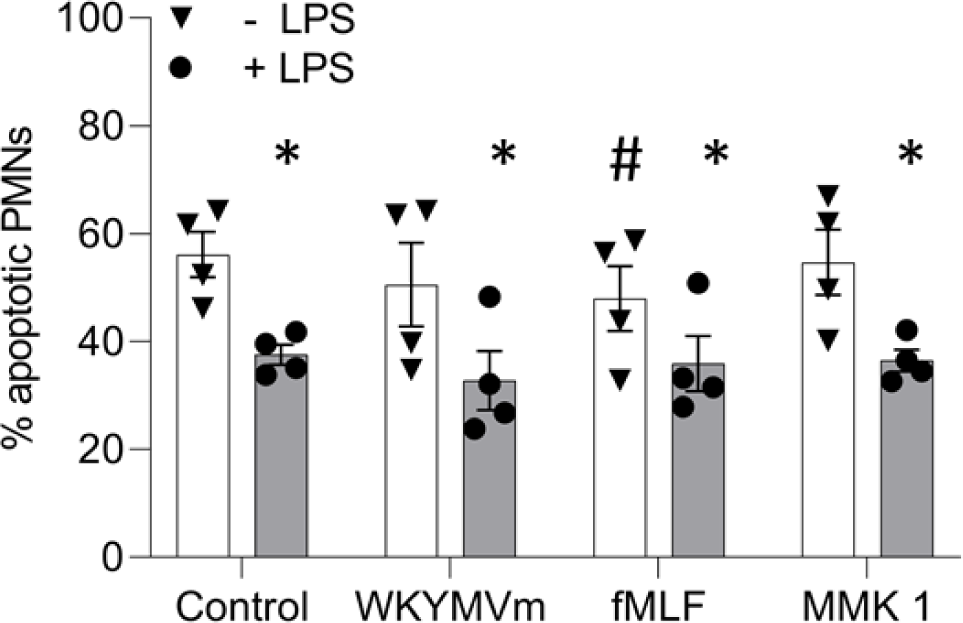
Survival of human neutrophils. Neutrophils were incubated with agonists (10-6 M) for 20 hours, with or without LPS (100 ng/mL). Bars display the mean ± SEM percentage of AnxA5+/PI-apoptotic human neutrophils of 4 individual donors represented by symbols. Repeated measurements One-Way ANOVA with Dunnett’s test. * p < 0.05, ** all vs ctrl; # p < 0.05 all vs ctrl within Group(-LPS/ +LPS).

## Discussion

Both FPR1 and FPR2 are crucial to infection and inflammation [13]. As the receptors share a high degree of similarity (69% amino acid sequence identity (Fig.1), it is thereby not surprising that they have an overlapping spectrum of ligands [6], [36].

Despite their similarity and shared agonists, they are believed to exert opposing functions. FPR1 mediates a wide array of pro-inflammatory responses [56]. In contrast, FPR2, coupled with its endogenous agonists, has been associated with agonist-induced anti-inflammatory and pro-resolving properties [26]–[28], [31], [57]. Studies have highlighted the pro-resolving role of FPR2 in cells lacking FPR2 functionality or when treated with annexin A and the annexin A1 pharmacophore peptide Ac2-26 which is an agonist for all receptors of the FPR family [50], [58]–[61].

Other endogenous FPR agonists encompass the neuroprotective humanin, which is encoded by a small 75-nucleotide open reading frame (ORF) located within the mitochondrial 16S rRNA gene [48], [62], [63]. However, several nuclear copies of humanin were identified. Therefore whether the corresponding RNA template is transcribed within mitochondria and translated by the mitochondrial ribosome or instead within the cytoplasm is yet to be conclusively established [64].

Meanwhile, other small peptides originating from the mitochondrial 16S rRNA gene have surfaced. Among these is the small humanin-like peptide 6 (SHLP6), adding to the pool of novel mitochondria-derived peptides with presumed functions in the regulation of apoptosis of cancer cells [45], [65], [66]. Considering the potential involvement in aging and inflammation, we successfully identified SHLP6 in our analysis as a new peptide agonist for both, FPR1 and FPR2. Noteworthy, humanin and SHLP6 analogs utilized here were non-formylated.

Recently, the interactions of various peptide ligands at both FPRs were resolved at the atomic level and broadened our understanding of FPR activation [67], [68]. Nevertheless, the systematic pharmacological comparison of peptide agonists across both receptors is lacking. Given the overall importance of these PRRs in inflammation and their potential as a druggable target to regulate inflammatory conditions, we set out to describe the FPR signaling repertoires. We focused on three typical pathways downstream of GPCRs that also have been linked with inflammation and are considered drug targets [69]–[71]. We chose the synthetic peptide agonist W-peptide, as the maximum activator at both FPR1 and FPR2 in our setting, as an ideal reference agonist for comparison within and also between receptors.

Despite differences in logistic parameters, both FPR1 and FPR2 exhibited a shared and conserved signaling profile across all agonists, showcasing a consistent pattern in their response and stressing the absence of origin-dependent agonist signaling textures. Both receptors seem inherently predisposed or “hardwired” toward a classical Gαi-mediated GPCR signaling profile including the inhibition of *de novo* cAMP formation, ERK1/2 activation, and receptor internalization [69]–[71]. However, individual ligands displayed preferences for specific pathways depending on the receptor. Therefore, the signaling outcome was defined by the specific pairing of agonist and receptor rather than being linked to a particular receptor isoform, signaling texture, or a specific class of agonists, such as formylated peptides. Even the anti-inflammatory peptide Ac2-26 [49] induced a similar signaling pattern at both receptors. In our analysis, we therefore did not identify a receptor-specific signaling texture that could discriminate receptor functions in the immune response.

Notably, FPR1 agonists were able to elicit maximum receptor activation across all pathways. On the contrary, there were no maximum activators at FPR2, neither formylated peptides nor non-formylated peptides, or proteins. Collectively, the existence of physiological maximum-activating agonists substantially distinguishes FPR1 from FPR2. This could be crucial for an orchestrated innate immune response: FPR1 agonists activate the full FPR1 response while FPR2 signals in scenarios involving substantial tissue degradation and the concomitant release of mitochondrial peptides.

We extended our investigation beyond *in vitro* studies to a classic immune cell model involving neutrophils [39]. Neutrophils, cells with abundant FPR expression, are among the first responders to inflammation, and their lifespan can dynamically change in response to inflammatory signals. Neutrophil-mediated host defense is increased through the prolonged neutrophil survival of these otherwise short-lived non-dividing cells [39], [72]. Because the resolution of inflammation involves the removal of activated neutrophils from the site of inflammation, enhanced neutrophil apoptosis is observed in the resolution phase while imbalanced neutrophil survival might perpetuate inflammation and tissue damage [73], [74]. Indeed, prolonged neutrophil lifespan is seen in chronic inflammation. Thus, neutrophil lifespan reflects the effectiveness of immune modulators [40]–[42]. Selective FPR1 activation by fMLF significantly lowered the number of apoptotic neutrophils, implying that the FPR1 signaling axis might influence neutrophil survival in a pro-inflammatory fashion. Surprisingly, FPR2-specific activation by MMK 1 did not significantly impact neutrophil survival nor was LPS-induced prolonged lifespan reversed; suggesting that the reported FPR2 pro-resolving function is not a property of the receptor but depends on a specific receptor-agonist combination.

## Materials and Methods

### Reagents

The adenylyl cyclase activator forskolin and the Gαi Inhibitor Pertussis Toxin were purchased from Tocris. The FPR-independent MAPKinase activator PMA was obtained from Sigma Aldrich. Peptide ligands were obtained and dissolved as listed.

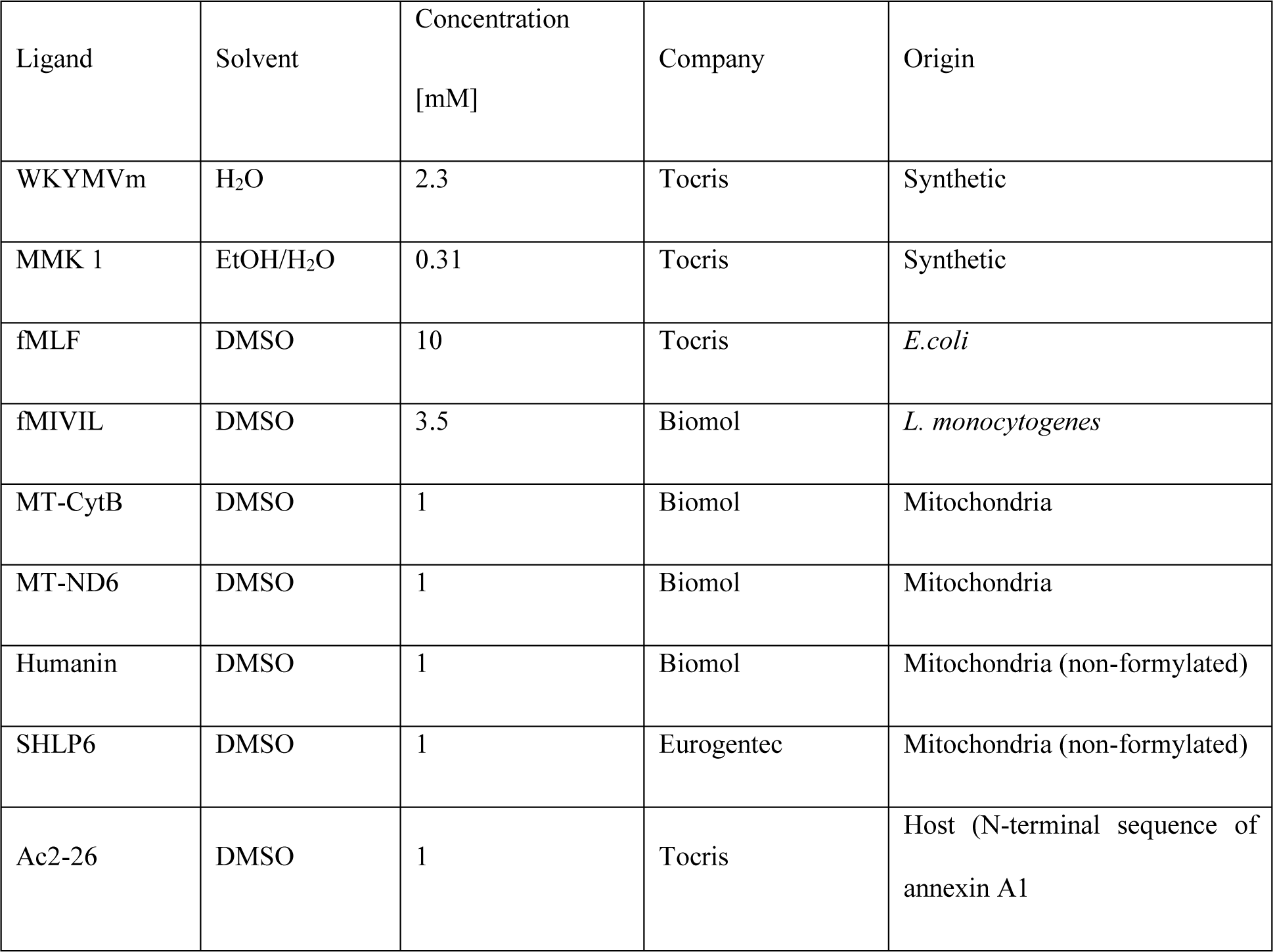

### Cell lines and culture conditions

The HEK293 FPR stable cell lines were generated as previously described [37]. In brief, cells were transfected with Lipofectamine 3000 with pcDNA3.1 (-) expression vector (Invitrogen, USA) encoding either FPR1 or FPR2 fused to a FLAG-tag.

HEK293 cell lines were maintained in Dulbecco’s Modified Eagle’s medium (DMEM, PAN Biotech, Germany) supplemented with 10% standardized fetal bovine serum (FBS Advanced, Capricorn-Scientific, Germany), 100 U/mL penicillin/0.1 mg/mL streptomycin (GE Healthcare, USA), 1% L-glutamine, 1% non-essential amino acids (NEAA, Sigma) at 37 °C and 7% CO_2_. Cell lines were routinely tested for the absence of mycoplasma contamination.

### Isolation of peripheral blood neutrophils

Human peripheral blood was donated by healthy volunteers upon informed consent. Blood drawing was approved by the institutional review board. Primary polymorphonuclear neutrophils (PMNs) were isolated with polymorphprep™ from Progen according to the manufacturer’s protocol. In brief, 1 volume of blood was added to 1 volume of pre-warmed polymorphprep and was centrifuged for 35 minutes at 500 x g (no breaks) at room temperature (RT). The PMN phase was collected and washed with PBS by centrifugation (5 minutes, 300 x g, 4 °C). The supernatant was discarded and erythrocytes were lysed by incubation with 1xRBC Lysis buffer (BioLegend). Neutrophils were resuspended and stored in Hanks balanced salt solution (HBSS) with Ca^2+^, and Mg^2+^, supplemented with 10 mM HEPES, 0.1% glucose, and 0.25% BSA, pH 7.4.

### Ex vivo assessment of human neutrophil survival

The method for the Investigation of prolonged neutrophil survival was adapted from Cho *et.al* [42]. Briefly, after neutrophil isolation and purification, 1 x 10^6^ cells were seeded in a 12-well suspension plate (Greiner bio-one) and incubated with RPMI 1640 VLE medium (Sigma Aldrich) at 37°C and 5% CO_2_ (untreated control). Treatment was performed by supplementing with LPS (100 ng/ml, Sigma-Aldrich, USA) and or indicated FPR ligands to the medium mentioned above. After 20 h of incubation, cells were washed twice with PBS (Pan Biotech, Germany) and labeled with propidium iodide (PI) and Annexin-V-FITC (Annexin V FITC Apoptosis Detection Kit with PI, Biolegend) for the detection of necrosis and apoptosis, whereas double negative cells were considered vital. The acquisition of 30000 cells was performed with a flow cytometer (Guava EasyCyte™, Merck Millipore) and data were analyzed using Incyte™ (Millipore).

### Analysis of gene expression via quantitative real-time PCR

Total RNA was isolated using the Trizol reagent (Invitrogen, Germany) according to the manufacturer’s protocol, and 1 µg RNA was reverse-transcribed using the High-Capacity cDNA Reverse Transcription Kit (Thermo Fisher Scientific, Germany). Gene expression levels were analyzed using predesigned QuantiTect primer assays (Hs_GAPDH_1_SG: QT00079247 Qiagen, Hs_FPR1_1_SG: QT00199745, Hs_FPR2_1_SG: QT00204295, Hs_FPR3_1_SG: QT00054677) and the Brilliant III Ultra-Fast SYBR Green qPCR Master Mix (Agilent Technologies, USA) on the Roche LightCycler480 PCR system (1 min 95 °C; 50 cycles of 5 sec at 95 °C, 10 sec 60 °C and final melting curve at 95 °C). Per cell line, at least three individual samples were run in triplicates.

### Analysis of HEK293-FPR cell surface expression via fluorescence-activated cell sorting (FACS)

The mouse monoclonal anti-FLAG antibody M1 (Sigma Aldrich) was labeled utilizing the DyLight488 Antibody Labeling Kit (Thermo Fisher Scientific) according to the manufacturer’s protocol. To determine the FPR cell surface pool and agonist-induced receptor internalization, *c*ells were cultured in 12-well flat-bottom plates and treated with vehicle or agonists diluted in DMEM for 20 min at 37 °C, and the reaction was subsequently stopped via cooling down on ice. Samples were transferred onto a 96-well v-bottom plate, washed two times with cell wash (PBS++/5%BSA/1mM CaCl_2_), and incubated with 5 µg/mL of the DyLight488-conjugated anti-FLAG M1 antibody for 45 min on ice. Surface was calculated as the difference between signals detected in vehicle-treated controls and the signals detected in agonist-treated cells. Beforehand, samples were incubated with 5 µg/mL of the DyLight488-conjugated anti-FLAG M1 antibody and the viability dye 7-Aminoactinomycin D (7-AAD, Invitrogen) for 45 min on ice and were then analyzed on a Guava easyCyte cytometer (Millipore). Median Fluorescence Intensity (MFI) values were determined for 10.000 488^+^/7-AAD^-^ cells.

### Analysis of de novo cAMP formation

FPR-mediated inhibition of cellular *de novo* cAMP formation was quantified by competitive immunoassays based on the time-resolved measurement of fluorescence resonance energy transfer (HTRF, Homogeneous Time-Resolved Fluorescence) using the cAMP-Gi Kit from Cisbio as previously published [37]. In brief, cells cultured in 96-well plates (50000 cells/well) were starved for 30 min in DMEM supplemented with 500 µM IBMX (Sigma Aldrich). To induce *de novo* synthesis of cAMP, cells were treated with forskolin (5 µM, Sigma Aldrich). Subsequently, FPR agonists at the indicated concentrations were added for 30 min at 37 °C, followed by cell lysis. Lysates were transferred onto a white opaque 384-well plate (Greiner Bio-one), the detection reagents (d2-labeled cAMP and anti-cAMP cryptate) were added, and samples were analyzed on the CLARIOstar reader (BMG Labtech) (200 flashes/well, integration start 60 µsec, integration time 400 µsec, settling time 100 µsec). Luminescence signals were expressed as the ratio 620/665 nm x the factor 10.000.

### Analysis of ERK1/2 activation

ERK1/2 phosphorylation in HEK293 FPR cell systems was measured with the HTRF immunoassay from Cisbio, as described previously [37]. In brief, cells seeded on a 96-well plate (30000 cells/well/25µL) in DMEM (without supplements) for 2 h were stimulated with agonists at the indicated concentrations for 5 min at 37 °C, followed by cell lysis for 30 min. After transferring the lysates to a 384-well low-volume plate, the labeled conjugates were added and the plate was incubated for 4 h at RT in the dark. The luminescence was recorded on the CLARIOstar reader (BMG Labtech) and the signals were expressed as the ratio of 10.000 x (acceptor signal/donor signal).

### Data analysis for pharmacological evaluation of agonist-mediated signaling profiles

For curve fitting, data of each assay were normalized as a fraction of the corresponding baseline (no agonist) and maximum system response. Datasets were analyzed using the GraphPad prism 10 built-in four parameters logistic model with the Hill slope set to 1. All data were tested for outliers (ROUT test method; Q value: 1%), which were excluded from further analysis. Data points represent the means ± standard error of the mean (SEM) of at least five independent experiments. The activity ratio logR (R = Emax/EC_50_) was calculated for each agonist and pathway. To evaluate the pathway preference of an agonist, i.e., the differential activities across pathways, ΔlogR_pathway_ values were calculated by subtracting the pathway1 logR values from the pathway2 logR values of the same agonist. For ranking the agonist activities within a pathway, the strong agonist W-peptide was chosen as a reference agonist and the W-peptide logR value was subtracted from each agonist’s logR value to generate ΔlogR_agonist activity_ values. All calculations were done according to the community guidelines for GPCR ligand bias [75].

### Statistical analysis

Results collected from the indicated biological replicates (n) are presented as mean ± SEM. Statistical analysis was performed using GraphPad Prism 8-10 (GraphPad Software, San Diego, California, USA). The normality of the data was analyzed by the Shapiro-Wilk test with α set at 0.05. The significance of the difference between two or more groups was analyzed by (repeated measurements) one-way ANOVA followed by Dunnett’s multiple comparisons tests, p < 0.05 (indicated by *) was considered significant. Actual p values are stated in the tables in suppl. Fig. S3.

## Author Contributions

DP and MFS performed the experiments and collected the data. OS and MB provided essential reagents and tools. DP, CAR, and UR designed and supervised the research analyzed the data, and wrote the paper. All authors revised the final version of the manuscript.

## Acknowledgments

This work is funded by the Deutsche Forschungsgemeinschaft (DFG, German Research Foundation) – CRC1009” Breaking Barriers”, project A06 to U.R. O.S. receives funding from the Deutsche Forschungsgemeinschaft (CRC TRR332 project A2, CRC1009 project B13, CRC1123 project A6, CRU342 project 1), Novo Nordisk, the Leducy Foundation, and the IZKF Münster. M.B. and U.R. are also supported by the Graduate School of Natural Products (GS-NP) of Münster University.

## Supplemental figures

**Suppl. Figure S1.**
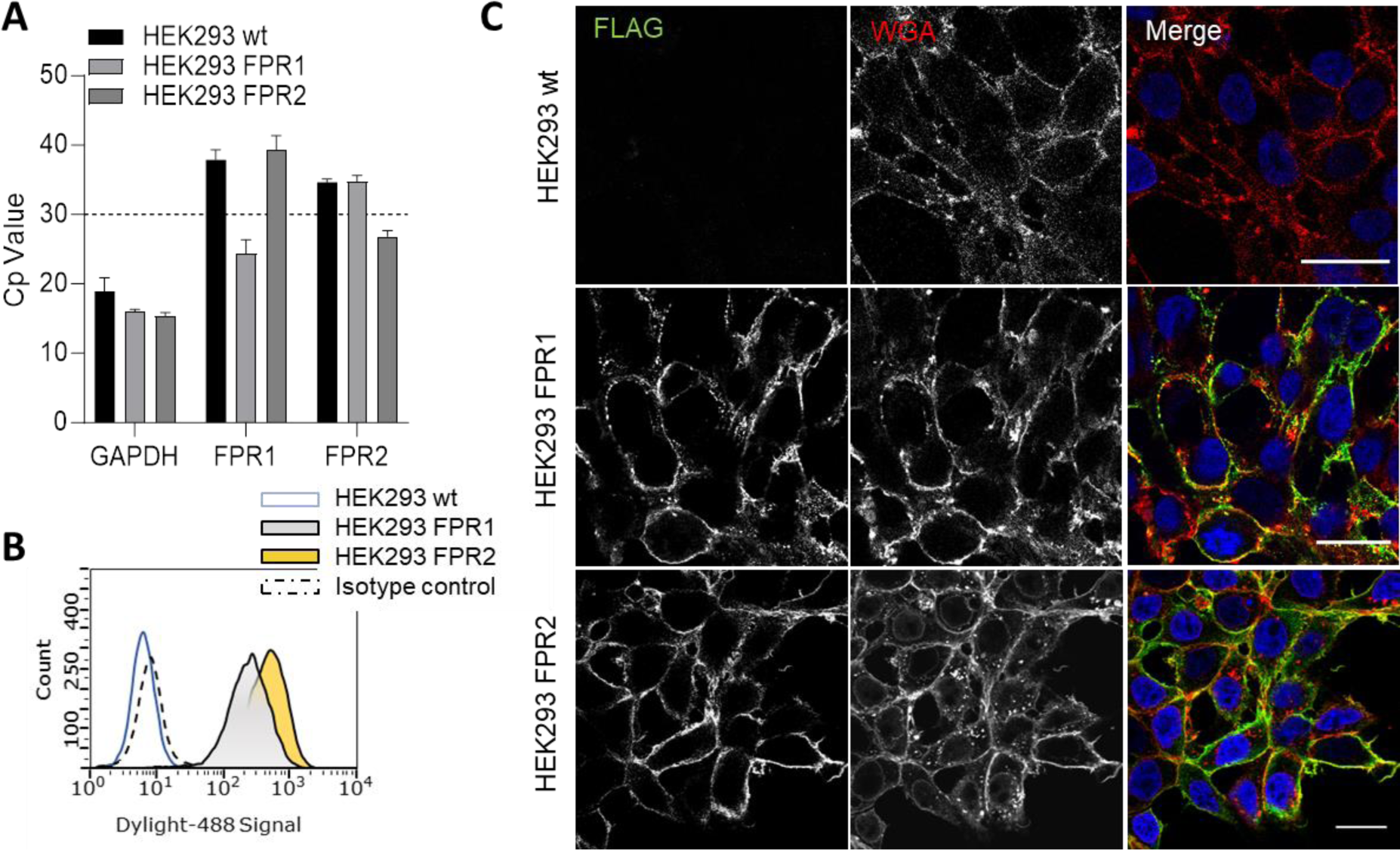
Characterization of the heterologous expression systems HEK293 FPR1 and HEK293 FPR2. (A) Gene expression analysis by qPCR confirming the absence of FPR1 and FPR2 expression in the parental cell line (wt) and validating the expression of the stably transfected FPR1 and FPR2 in the corresponding cell lines. Glycerinaldehyde-3-phosphate-Dehydrogenase (GAPDH) served as the reference gene. The threshold cycle value (Ct) considered relevant for protein expression was set to 30 (dotted line). Data are presented as the mean ± SEM (n = 4). B) Flow cytometric analysis of cell surface receptor expression quantified as a shift in mean fluorescence intensity versus parental HEK (wt). Cell surface-associated FLAG epitopes were detected with Dylight-488-coupled anti-FLAG antibody. The dotted line depicts the isotype control, n = 6. (C) Immunofluorescence analysis confirms the correct plasma membrane localization of the FLAG-tagged receptors. Fixed cells were stained with an anti-FLAG antibody (green), and the plasma membrane was visualized with wheat germ agglutinin (WGA, red). Nuclei were stained with DAPI (blue). A representative confocal image is shown. Scale bar, 20 µm.

**Suppl. Figure S2.**
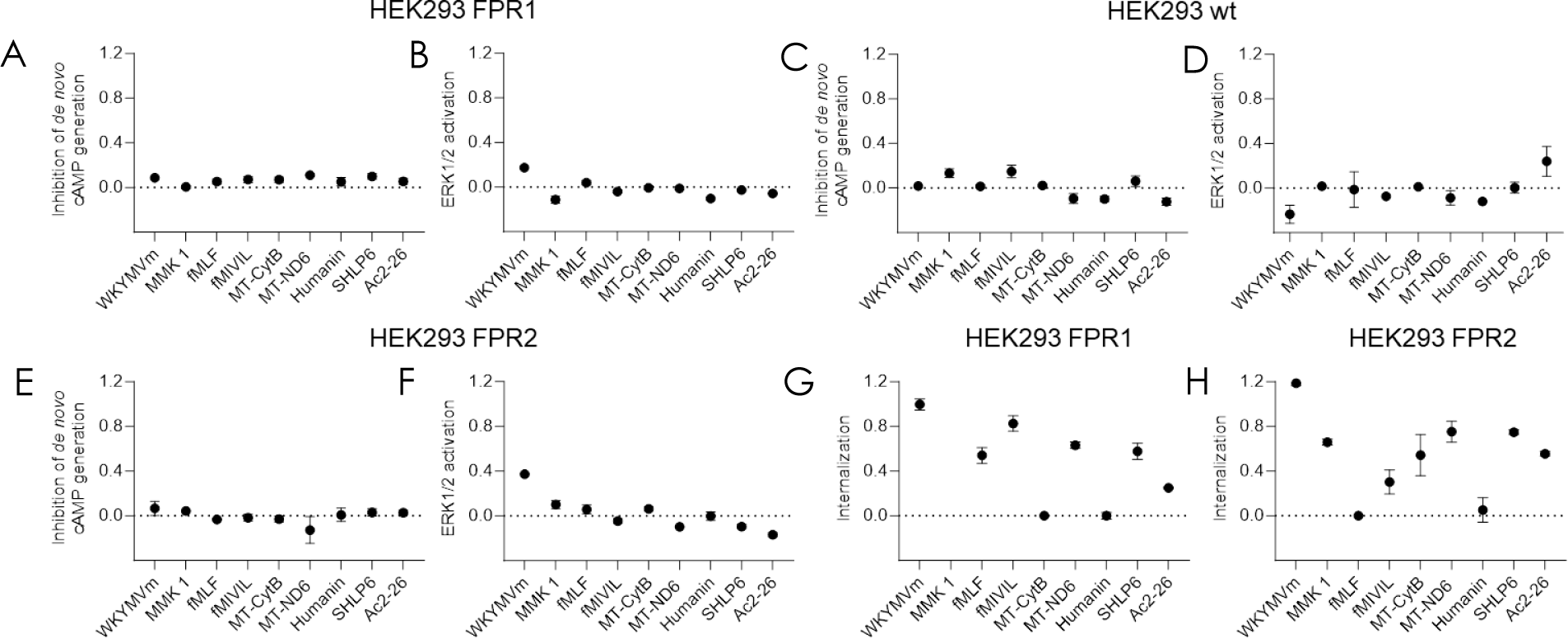
Analysis of G protein dependency. HEK293 FPR1 and HEK293 FPR2 cells were incubated with the indicated agonists (10^-6^ M) in the presence of pertussis toxin and responses were measured for A/E) the inhibition of cAMP synthesis, B/F) ERK1/2 activation and receptor Internalisation (G/H). (C/D) HEK293 wt cells were treated as in A and B in the absence of PTX. Data points display the mean ± SEM response for each pathway of n = 5 independent experiment.

**Suppl. Figure S3.**
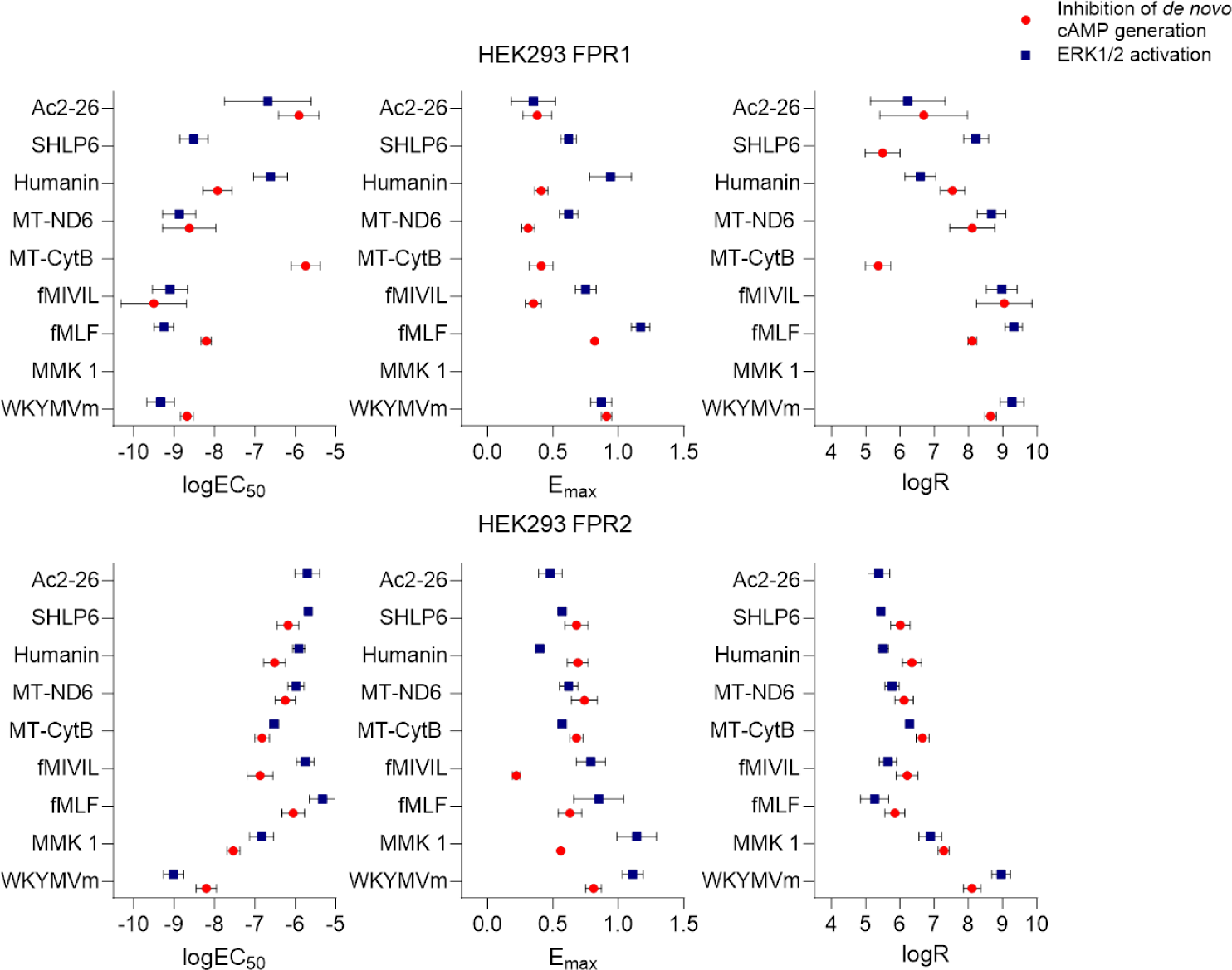
Overview of the logistic parameters for FPR1 and FPR2 across pathways. To comprehensively evaluate agonist performance, we calculated logR values, which integrate both logEC50 and the Emax of the respective agonist, for each pathway.

**Suppl. Figure S4.**
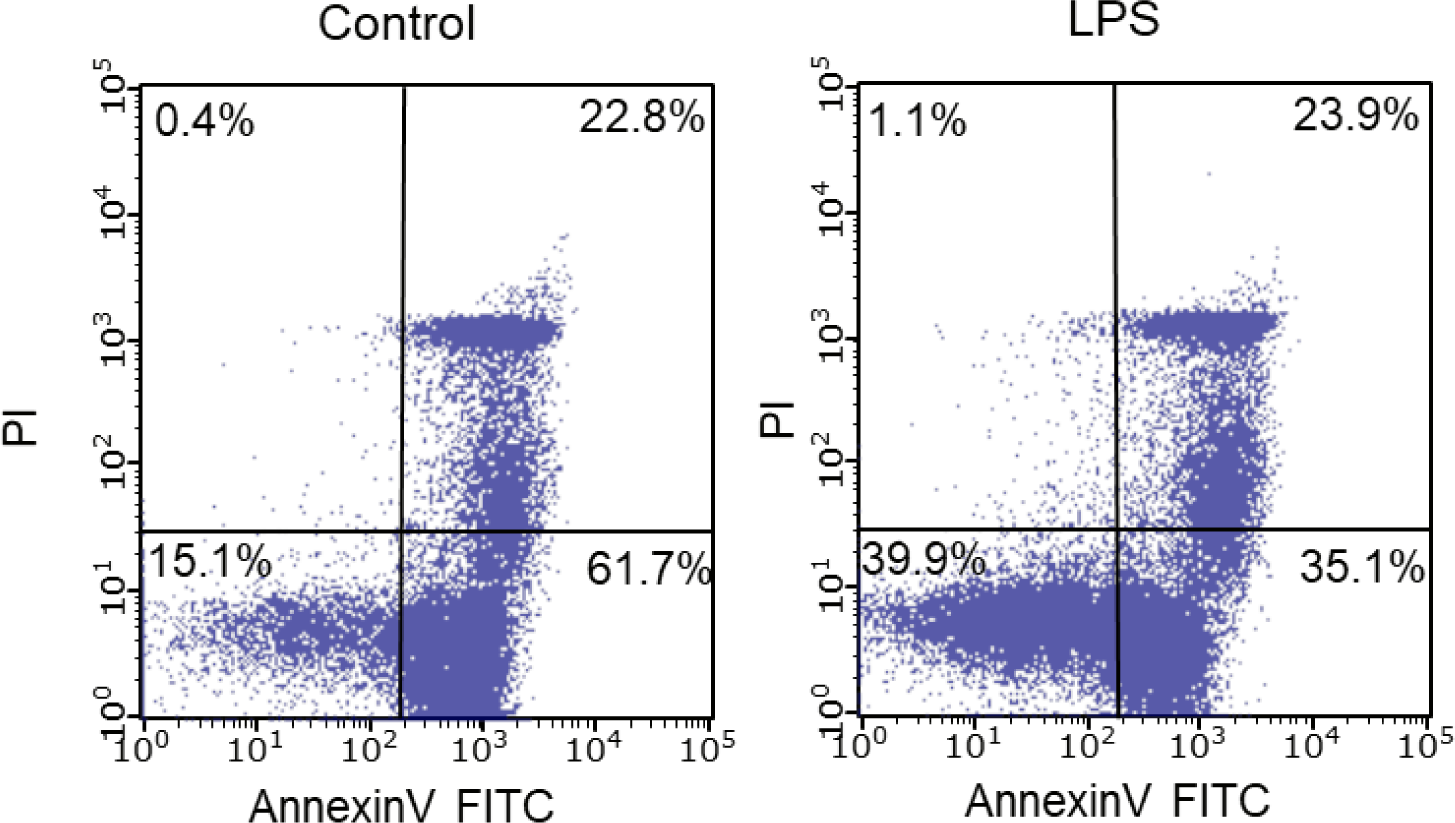
LPS-induced prolonged Neutrophil survival. Isolated neutrophils were treated with LPS (100ng/ml) for 20 hours, or left untreated, and subsequently stained with propidium iodide (PI) and FIC -coupled AnnexinV to determine cell survival state. Sown are representative original dot blots of untreated and LPS-treated neutrophils of a single representative donor.

**Suppl. Table I.**
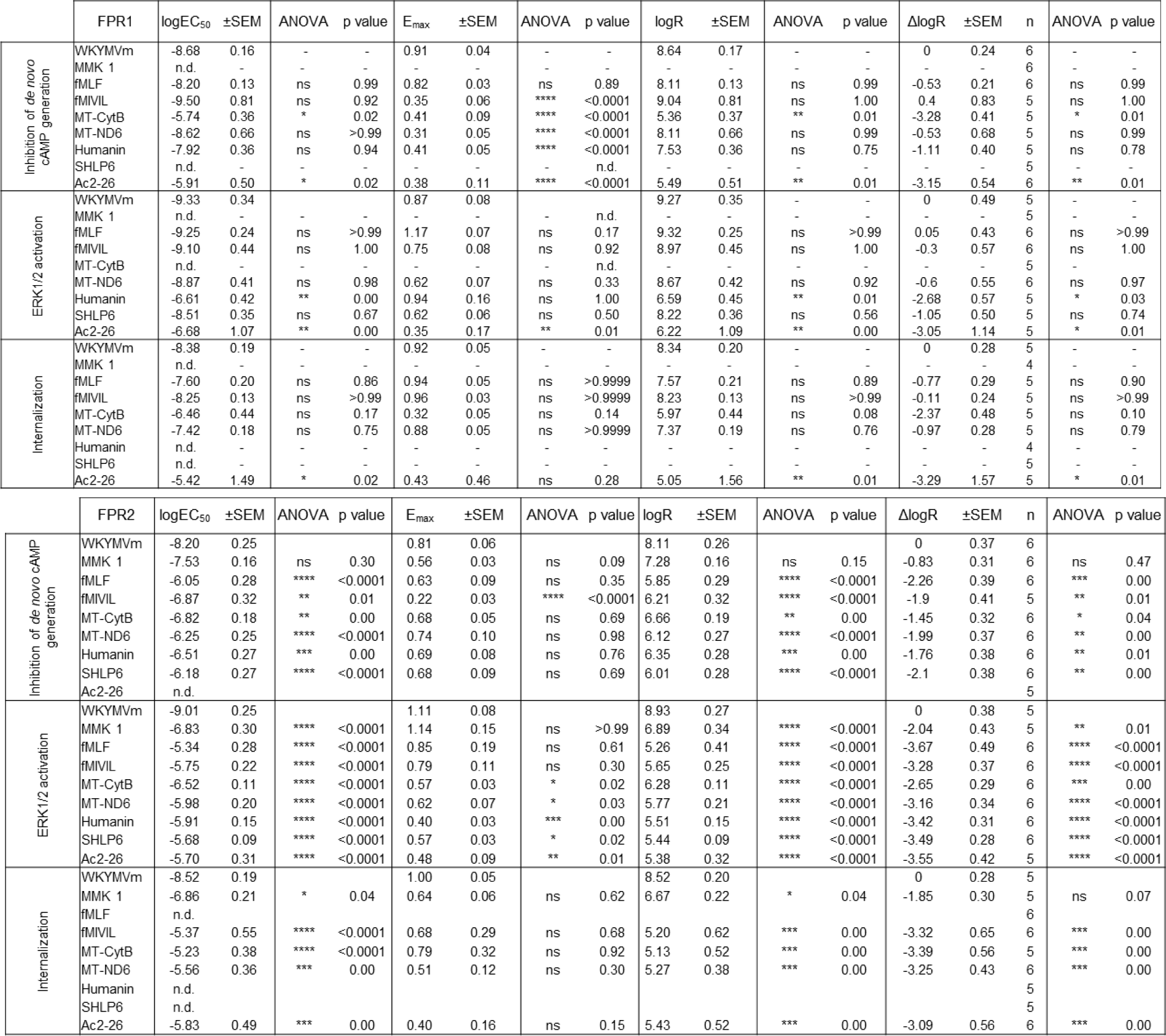
Effects of FPR ligands in cAMP inhibition, ERK1/2 activation, and receptor internalization experiments performed in HEK293 cells expressing FPR1 or FPR2. The table lists the concentration-response curves-deduced pharmacological parameters, logEC_50_ and Emax as well as the calculated parameters logR (activity ratio) and ΔlogR. The table provides corresponding p values and significance indicators according to ANOVA followed by the Dunnett test. Data are mean values of at least 4 experiments. N.d. = not detectable; ns = not significant.

## Notes

### Competing Interest Statement

The authors have declared no competing interest.

